# Limited differential expression of miRNAs and other small RNAs in LPS-stimulated human monocytes

**DOI:** 10.1101/461277

**Authors:** Daniel Lu, Tracy Yamawaki, Hong Zhou, Wen-Yu Chou, Mark Chhoa, Edwin lamas, Sabine S. Escobar, Heather A. Arnett, Huanying Ge, Songli Wang, Chi-Ming Li

**Author notes:** To whom correspondence should be addressed: Chi-Ming Li PhD, 1120 Veterans Blvd, MS: ASF1-2, South San Francisco, CA94080, E-Mail, (Office).

## Abstract

Monocytes are a distinct subset of myeloid cells with diverse functions in early inflammatory immune modulation. While previous studies have surveyed the role of microRNA regulation on different myeloid cell lines and primary cultures, the time-dependent kinetics of inflammatory stimulation on miRNA expression and the relationship between miRNA-to-target-RNA expression have not been comprehensively profiled in monocytes. In this study, we use next-generation sequencing and RT-PCR assays to analyze the non-coding small RNA transcriptome of unstimulated and lipopolysaccharide (LPS)-stimulated monocytes at 6 and 24 hours. We identified a miRNA signature consisting of five mature miRNAs (*hsa-mir-146a*, *hsa-mir-155*, *hsa-mir-9*, *hsa-mir-147b*, and *hsa-mir-193a*) upregulated by LPS-stimulated monocytes after 6 hours and found that most miRNAs were also upregulated after 24 hours of stimulation. Only one miRNA gene was down-regulated and no other small RNAs were found dysregulated in monocytes after LPS treatment. In addition, novel tRNA-derived fragments were also discovered in monocytes, although none showed significant changes upon LPS stimulation. Interrogation of validated miRNA targets by transcriptomic analysis revealed that the majority of differential mRNA expression was maintained in LPS-stimulated monocytes although few of miRNA targets seemed to obtain heterogeneous expression pattern along the treatment. Our findings reveal a potential role by which selective miRNA upregulation and stable expression of other small RNAs enable monocytes to develop finely tuned cellular responses during acute inflammation.

## Background

Monocytes belong to a subset of circulating white blood cells that can further differentiate into macrophages and dendritic cells in solid tissues surrounding sites of injury [1]. *In vivo* and *in vitro* studies have shown that monocytes and their derivatives function as an essential component of the innate immune system that mediate the host defense, serve as the first line of resistance to microbial attack, modulate tumor-associated immune defense responses, and regulate tissue homeostasis [2]. Indeed, the pleiotropic potential of monocytes suggests that their cellular functions must be tightly regulated in a manner unique from more differentiated myeloid cell types to enable appropriate, context-dependent responses.

Micro-RNAs (miRNAs) are a large class of small noncoding RNAs that post-transcriptionally regulate mRNAs and subsequently influence essential cellular functions through modulation of gene expression at the RNA or protein level [3]. miRNAs are initially transcribed as long primary transcripts (pri-miRNA) by mammalian type II RNA polymerases and then processed into precursor hairpin intermediates (pre-miRNA) in cell nuclei by an RNase III (Drosha). After transport to cytoplasm by an export complex, Exportin5-RanGTP, pre-miRNAs are further processed by another cytoplasmic RNase III, Dicer, to generate 18–24 nucleotide mature miRNAs through a highly regulated biogenesis process [4]. Mature miRNAs are then loaded into the RNA induced silencing complex (RISC) which targets specific mRNA causing translational repression or degradation of the targeted mRNAs [5]. Each miRNA can potentially target dozens to hundreds of mRNAs, and, thus, miRNAs have been implicated in regulating multiple and disparate biological pathways [4–6].

Several miRNAs in myeloid cells have previously been identified that impact immune responses, although results have varied due to different cell types, assay methods, and culture/stimulation conditions [7,8] (summarized in Supplementary Table 1). Expression analyses using oligo-based microarray technology revealed that *hsa-mir-155* was highly induced during the maturation of dendritic cells derived from human monocytes and in monocyte cell lines [9,10]. This finding was further confirmed with the identification of other miRNAs, including *hsa-mir-9* and *hsa-mir-146a*, which were up-regulated in LPS-treated monocytes using a qPCR-based platform [11]. In each study, however, a genome-wide assessment of small RNA expression profiles was limited due to the use of oligo-based or qPCR-based assay platforms. Next-generation sequencing (NGS) technologies have significantly increased the ability to detect low abundant and novel small RNAs in numerous human cell types [12–14]. This method enables the isolation and detection of all small RNAs without *a priori* knowledge or bias, allowing for global expression profiling of small RNAs relevant to activated monocytes. Such analyses have also led to the discovery of small RNA species other than miRNAs that include RNA fragments derived from small nucleolar RNA (snoRNA) and transfer RNA (tRNA) [15]. While the function of these snoRNA-derived (sdRNA) and tRNA-derived (tRF) RNAs is obscure, their expression is altered in certain human diseases [15–17]. Further analyses of the functional role of smRNAs and tRFs may lead to the development of disease-associated biomarkers or avenues for novel therapeutic strategies.

In this study, we performed deep sequencing to interrogate the expression of small RNAs in non-stimulated and LPS-stimulated human monocytes from multiple donors. The small RNA transcriptome mainly consisted of miRNAs but also contained snoRNAs, rRNAs, Y RNAs, and tRNAs. We find that a small set of miRNAs are stably upregulated following LPS stimulation at two different timepoints and confirm these findings using multiple RT-PCR assays. We also examine how miRNA upregulation affects target mRNA levels by analyzing the RNA expression of stimulated monocytes using RNA-seq. Furthermore, we describe, for the first time, identification of 18 tRNA-derived fragment RNAs in monocytes. Further analyses of these identified small RNAs may lead to a better understanding of their role in monocyte biology and serve as biomarkers to detect monocyte function in human disease.

## MATERIAL AND METHODS

### Preparation and lipopolysaccharide (LPS) treatment of human monocytes

#### Ethics Statement

Samples were collected anonymously through an internal blood donor program that was monitored by the Western IRB Board and was considered exempt from IRB approval.

Primary human monocytes were freshly isolated from peripheral blood mononuclear cells (PBMCs) from individual donors using the EasySep Monocyte Enrichment kit from Stem Cell Technologies (negative selection). Cells underwent analysis by flow cytometry for purity (CD14-PE) (Becton Dickerson), with donor preps having cell purity greater than 90% for deep sequencing and further validation. Monocytes were suspended at 10 million/ml in X-Vivo medium (Lonza) supplemented with 5% AB human serum (Invitrogen) and incubated for 6 or 24 hours with either medium only or with media containing 100 ng/ml lipopolysaccharide (InVivoGen). All cells (both adherent and suspension) were harvested, washed, and pelleted, with the cell pellet snap-frozen on dry ice and placed at −80°C until further analysis.

### RNA isolation

Cell pellets containing 10-20 million monocytes each were lysed and homogenized in 1ml of the Trizol reagent from Life Technologies. Following commercial instructions, sedimentation of the aqueous and organic phases was carried out by vigorous shaking after adding 0.2 ml chloroform and subsequent centrifugation of the samples at 14,000 RPM for 20 minutes using a bench top centrifuge (Eppendorf). The aqueous phase was carefully transferred to a new 1.5ml tube and then mixed with 0.45 ml of 100% isopropanol. The RNA was centrifuged and precipitated at 14,000 RPM for 20 minutes. After washing with 75% ethanol, the RNA was dissolved in DEPC-treated water and its quality and concentration were determined using Bioanalyzer (Agilent) and Nanodrop (Thermo Scientific), respectively.

### Small RNA library Construction

Barcoded cDNA Libraries were generated from 1ug of total RNA from each human monocyte sample using the Illumina TruSeq small RNA kit per the manufacturer’s protocol. The small RNA libraries were further purified by electrophoresis in a 6% PAGE gel. After extraction from the gel slices between 140-160 bp, the libraries were concentrated by ethanol precipitation. cDNA libraries were suspended in 10mM Tris, pH 8.0 buffer and the DNA concentration was determined using Taqman-based qPCR assay.

### Deep sequencing and data analysis of small RNA libraries by Illumina NGS

Approximately 1.2 pmol of small RNA libraries were pooled and sequenced using either a HiSeq or GA flowcell. Clusters were generated using an Illumina C-BOT instrument. One multiplexing single read, 30/7, was set up and performed on the Illumina HiSeq 2500 or Illumina GAIIx using the small RNA sequencing kit. Once FASTQ files were generated from raw BCL output, the total reads of each library were batched into individual bins based on sample barcode indices. The adaptor sequencing was stripped from each read, and reads without the 3′ adaptor sequence were filtered out. Reads with low quality (Phred Q<30) or length ≤12 nt after adaptor stripping were further removed. For the remaining high-quality reads, we counted the occurrence of each unique sequence in each sample and compressed the read files to contain only unique sequences. Unique reads were then aligned to small RNA reference library using the OmicSoft Aligner (V5.1)[18]. The small RNA reference library includes: miRNA hairpin sequences from miRBase (v20) [19], tRNA sequences from GtRNAdb [20,21], snoRNA/scaRNA sequences from snoRNABase [22], rRNA and Y RNA 1/3/4/5 from NCBI. After mapping, reads that aligned to a unique reference sequence were retained, and read abundances were normalized to counts per million (CPM). DESeq2 [23] was used to test for differential expression of small RNAs, controlling for the effect of the donor on normalized expression (*i.e.* design ~ donor + LPS.treatment).

### Expression Validation by Taqman-based qRT-PCR assays

1 ug of total RNA was treated with DNase 1 (New England Biolabs) in 50 ul 1x DNase1 Buffer as outlined in the commercial protocol. All the miRNA-specific and U6 Taqman-based assays and reagents were acquired from the commercial collection at ABI/Life Technologies. After reverse transcription with the primers specific for individual miRNAs that were provided in the assays, the transcribed cDNA was applied for quantification by using Taqman-based assays on ABI 7900HT. This was indicated as the relative expression to the level of U6 RNA according to ΔΔCt that was calculated and generated using the ABI SDS version 2.4 software after normalization of RNA input.

### Tagged RT-PCR Semi-Quantification for mature and precursor miRNAs

1 ug of DNase1-treated total RNA was then heated to 70°C for 2 minutes to eliminate secondary structure, and immediately cooled on ice. The 5’ and then the 3’ adapters were ligated to the sample using the miRCat33 (Integrated DNA Technologies; IDT) in subsequent reactions with a sodium acetate clean-up between each reaction as outlined in the protocol. These reactions were followed by reverse transcriptase using Invitrogen Superscript III, and then diluted 4x with IDTE (pH 7.5) supplied in the miRCaT33 Kit. The primers that were reversed and complimentary to the miRNA specific sequences were designed and ordered from IDT. These primers coupled with the miRCAT forward or reverse primers (depending on the orientation of the miRNA sequence on the precursor miRNA) were included in a PCR reaction using Promega GoTaq^®^ Hot Start Green Master Mix. Optimum cycles varied per miRNA based on the level of expression, typically starting at 25 cycles for *hsa-mir-155* and *hsa-mir-21*, 28 cycles for *hsa-mir-146a* and *hsa-mir-147b, and* 31 cycles for *hsa-mir-9* and *hsa-mir-193a* by using the following conditions: 1 cycle of 95°C for 3min and 30 seconds, 25-31 cycles of 95°C for 30 seconds, 52°C for 30 seconds, and 72°C for 40 seconds, 1 cycle of 72°C for 5 minutes, then a 4°C hold. The amplified products were then resolved by electrophoresis in 6% polyacrylamide or 4% EtBr-containing agarose gels along with the 25 bp DNA Ladder. To stain the polyacrylamide gels, the exposed gel was incubated in the GelStar Gel Stain (1:1000 dilutions) from Lonza for 15 minutes. The gels were then imaged using a digital camera over a UV transluminator from BioRad. The products were further validated by Sanger sequencing after the procedure of TA-cloning (Invitrogen). The details of the primers are available upon request.

### Deep sequencing and data analysis of mRNA libraries by Illumina NGS

2ug total RNA was used for cDNA library preparation by using a modified protocol based on the Illumina Truseq RNA Sample Preparation Kit V2. After poly-A selection, fragmentation, and priming, reverse transcription was carried out for 1^st^ strand cDNA synthesis in the presence of RNaseOut (Invitrogen) and actinomycin-D (MP Biomedicals). The synthesized cDNA was further purified by using AMPure RNAClean beads (Beckman Coulter). A modified method by incorporation of dUTP instead of dTTP was prepared and used for the second strand synthesis. After AMPure XP bead purification (Beckman Coulter), following the standard protocol recommended by the Illumina Truseq RNA kit, end repairing, A-tailing, and ligation of index adaptors were sequentially performed for generation of cDNA libraries. After size selection of libraries using Pippin Prep (SAGE Biosciences), the dUTP-containing strands were destroyed by digestion with USER enzymes (New England Biolabs) followed by PCR enrichment. Final cDNA libraries were analyzed in Agilent Bioanalyzer and quantified by Quant-iT^TM^ Pico-Green assays (Life Technologies) before sequencing using the HiSeq platform (Illumina).

### tRF validation by Stem loop RT-PCR

500ng DNase1-treated total RNA from each sample was incubated with 2 U RNase Inhibitor and 10 U Superscript II in the 1^st^ cDNA buffer containing 10mM DTT, 250 uM dNTP, and 50 nM stem loop RT primers for *tRF-GlnTTG*, 5’-gtcccagcaggtgcagggtccgaggtattcgcacctgctgggac<u>tcggatc</u>-3’, *tRF-SerTGA*, 5’-gtcccagcaggtgcagggtccgaggtattcgcacctgctgggac<u>aaaataag</u>-3’ and *tRF-TyrGTA*, 5’-gtcccagcaggtgcagggtccgaggtattcgcacctgctgggac<u>cttcgag</u>-3’. The reaction for stem loop reverse transcription started from 30 minutes of incubation at 16 °C followed by 60 cycles of 30°C for 30 seconds, 42°C for 30 seconds, and 50°C for one second and ended with one step of 85°C for 5 minutes. The tRF cDNAs were amplified with the forward primer specific for each tRF, 5’ - accacgactttgaatccag-3, 5’-accacgaagcgggtgct-3, or 5’-tcgtcctggttcgattcc-3’, and a common reverse primer, 5’-gcagggtccgaggtattc-3’ by using NEB phusion HF Taq and the following conditions: 1 cycle of 98°C for 30 seconds, 10 cycles of 98°C for 10 seconds, touch-down annealing from 64 to 55°C for 30 seconds, and 72°C for 30 seconds, 20 cycles of 98°C for 10 seconds, 55°C for 30 seconds, and 72°C for 30 seconds, and 1 cycle of 72°C for 5 minutes, then a 4°C hold. The amplified tRF products were further resolved in 1x TBE 6% polyacrylamide gels. After staining in the GelStar Gel Stain buffer, the gels were imaged using a digital camera over a UV transluminator from BioRad.

## RESULTS

### Deep sequencing of small RNAs from human monocytes

Deep sequencing for small RNAs was performed by constructing 18-30-bp cDNA libraries from human monocyte RNA samples that were collected from seven different healthy donors. These purified monocytes had been treated by LPS or control medium alone for either 6 or 24 hours to assess how the duration of stimulation affects miRNA expression. The resulting libraries were then sequenced using Illumina instruments to generate an average of 6.4±2.2 million sequencing reads per sample (Supplementary Table 2). An analysis pipeline elucidated in Supplementary Fig 1 was applied for data QC, read alignment, and small RNA transcriptome analysis. Individual sequences were aligned to a miRNA hairpin sequence database from miRBase and other small RNA reference databases (tRNA sequences from GtRNAdb; snoRNA/scaRNA sequences from snoRNABase; ribosomal RNAs and hY RNAs 1/3/4/5 from NCBI) using the OmicSoft Aligner.

An average of 54.0% of the unique reads that passed the QC metrics mapped uniquely to one of the small RNA references listed above (Supplementary Table 2). 72.1-95.8% of the mapped reads were specifically categorized to miRNA (Supplementary Fig 2), which spanned 20 to 24 bp in sequence length (Supplementary Fig 3). Among the 1,859 miRNA sequences listed in miRBase, 451 were detected in all samples while 850 miRNAs were detected in over half of the samples. While expression of miRNAs spanned five orders of magnitude, the top 100 expressed miRNAs comprised 92% of the entire miRNA population detected in human monocytes, revealing that miRNA gene diversity is largely restricted to the most highly expressed miRNAs (Supplementary Fig 4). The remainder of the mapped reads aligned to rRNAs (2.8-15.4%), snoRNAs (0.60-13.7%), tRNA (0.23-1.2%), and hY RNAs (0.41-7.3%) (Supplementary Fig 2).

### miRNA transcriptome analysis in human LPS-treated monocytes

To determine whether monocytes stimulated by LPS differentially express miRNAs relative to monocytes cultured only in medium, we performed differential expression testing between monocytes that were cultured for the same duration using DESeq2, adjusting for patient-specific effects and using normalized read counts. We found that 15 and 13 miRNAs are differentially expressed between LPS- and medium-treated monocytes at 6 and 24 hours, respectively, after correcting for multiple testing (adjusted *p-value* < 0.05, FDR=0.01) (Tables 1 and 2) (Fig 1A-1B). At 6 hours, all 15 miRNAs were upregulated with 1.7 – 11.2-fold upregulation (Fig 1A). At 24 hours, the magnitude of upregulation was more pronounced relative to the monocytes stimulated for 6 hours, with 12 out of the 13 miRNAs having 2.3 – 55.3-fold upregulation (Fig 1B). One miRNA, *hsa-mir-7151* was downregulated in monocytes stimulated with LPS for 24 hours. Five miRNA species (*hsa-mir-155*, *hsa-mir-146a*, *hsa-mir-9*, *hsa-mir-147b*, *hsa-mir-365b* and *hsa-mir-193a*) were upregulated at both timepoints from the NGS data and may serve as steady regulators of the inflammatory response of monocytes. Of the stably upregulated miRNAs, only *hsa-mir-155* and *hsa-mir-146a* were among the top 100 most highly expressed miRNAs (Supplementary Fig 4). Examination of each donor sample sequenced revealed high patient-to-patient variability in baseline expression of each miRNA, as the miRNA expression level in media-treated monocytes of some patients exceeded the expression level in LPS-treated monocytes of other patients (Fig 1C-1D). However, intra-patient differential miRNA expression was consistent after LPS stimulation for all differentially expressed miRNAs, underscoring the importance of correcting for patient variance and revealing that relative miRNA changes rather than absolute miRNA levels may be more important for the induction of monocyte responses to inflammatory stimuli.

**Figure 1.**
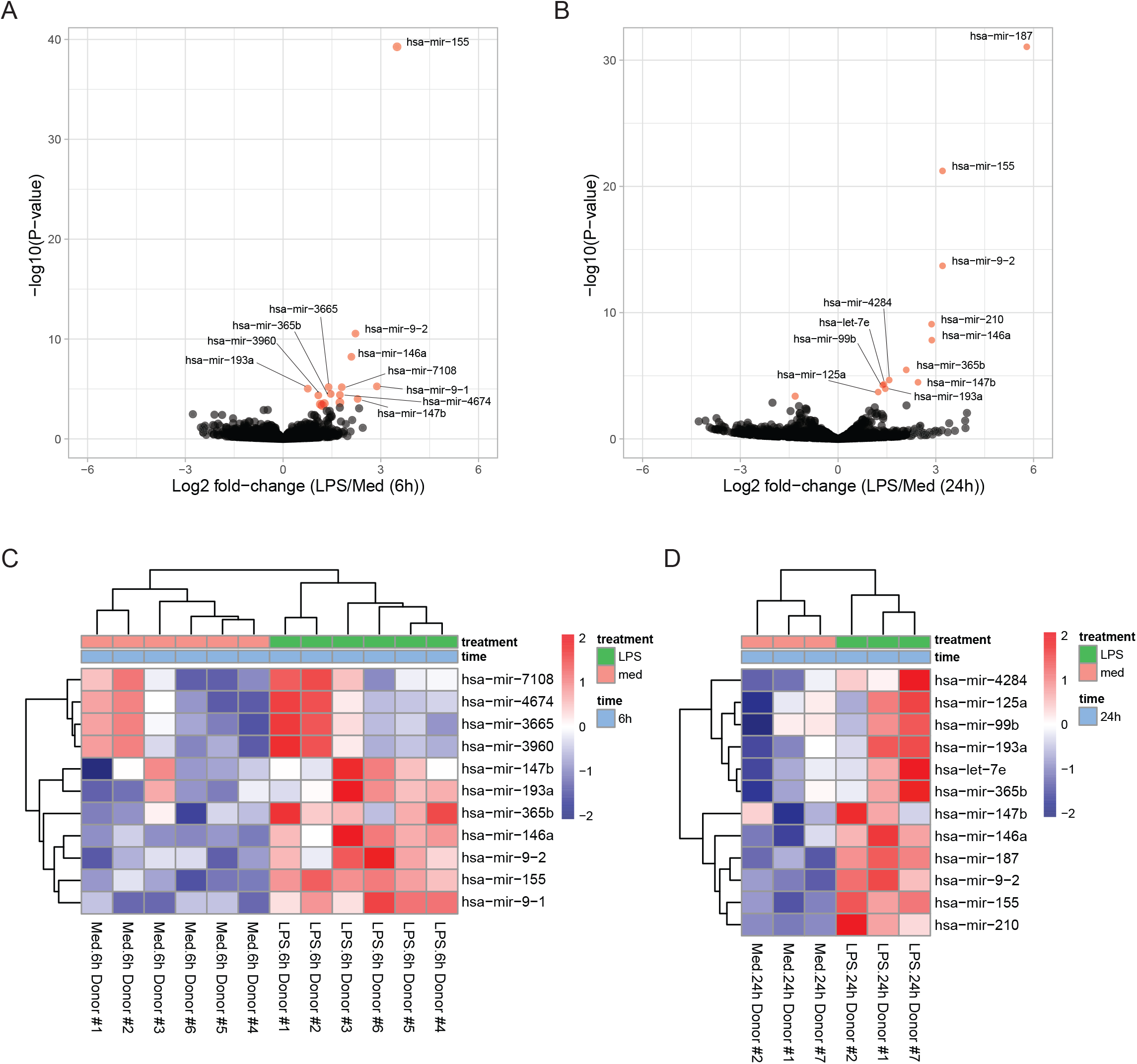
LPS consistently upregulates certain miRNAs in LPS-treated monocytes. (A and B) Differential expression analysis of the miRNA transcriptome is summarized as a volcano plot of the log2 fold-change in miRNA expression in LPS-stimulated monocytes versus medium-only-treated monocytes treated for either (A) 6 hours, or (B) 24 hours. miRNAs differentially expressed with an adjusted p-value<0.05 (FDR=0.01) are depicted in red. (C and D) Heatmap and hierarchical clustering depicting the normalized Z-scaled expression of each highly differentially expressed miRNA (adj.p-value<0.01) for each donor and sample at (C) 6 hours, or (D) 24 hours.

### Expression Validation of mature miRNAs by quantitative RT-PCR and pre-cursor miRNAs by tagged RT-PCR

To confirm upregulation of signature miRNAs that were observed by NGS in LPS-stimulated human monocytes, we utilized quantitative real-time PCR to analyze the expression patterns of miRNAs upregulated at both timepoints, one of the timepoints, and at neither timepoint. Of the miRNAs upregulated at both timepoints by NGS, we confirmed upregulation of *hsa-mir-155, hsa-mir-146a, hsa-mir-9*, and *hsa-mir-147b* (Fig 2A-2D). (*Hsa-mir-9-1* and *hsa-mir-9-2* both produce the same mature miRNA (*hsa-mir-9*) and are detected in the same RT-PCR reaction.) RT-qPCR also identified upregulation at 24 hours but not after 6 hours for *hsa-mir-187, hsa-mir-193a, hsa-mir-99b, hsa-mir-125-5p*, or *hsa-mir-125-3p* (Fig 2E-2I). This corroborated NGS results for all miRNAs except for *hsa-mir-193a*, which originally had shown the most modest upregulation (1.7-fold) out of the upregulated miRNAs in the NGS data (Fig 1A). *Hsa-mir-365b* showed upregulation at neither timepoint by qRT-PCR (Fig 2J). *Hsa-mir-146b, hsa-mir-149*, and *hsa-mir-2116* were also included as negative controls for qRT-PCR based on NGS results. *Hsa-mir-146b*, which was previously identified in another study [11], was selected as a negative control since it differs from *hsa-mir-146a* by two nucleotides. *Hsa-mir-449c, hsa-mir-149*, and *hsa-mir-2116* showed similar expression in both treatment groups. Importantly, none of the miRNAs selected as negative controls were differentially expressed by RT-qPCR (Fig 2J-2M). Overall, RT-qPCR confirmed time-dependent miRNA expression patterns for 25 of the 28 conditions tested across 14 miRNAs (*hsa-mir-193a* (Fig 2F) and *hsa-mir-365b* (Fig 2N) were the only miRNAs with discrepancies), demonstrating high concordance between the two methods. In correlation with the NGS results, upregulated miRNAs detected by qRT-PCR generally tended to have greater relative expression difference in LPS-stimulated monocytes at 24 hours compared to 6 hours.

**Figure 2.**
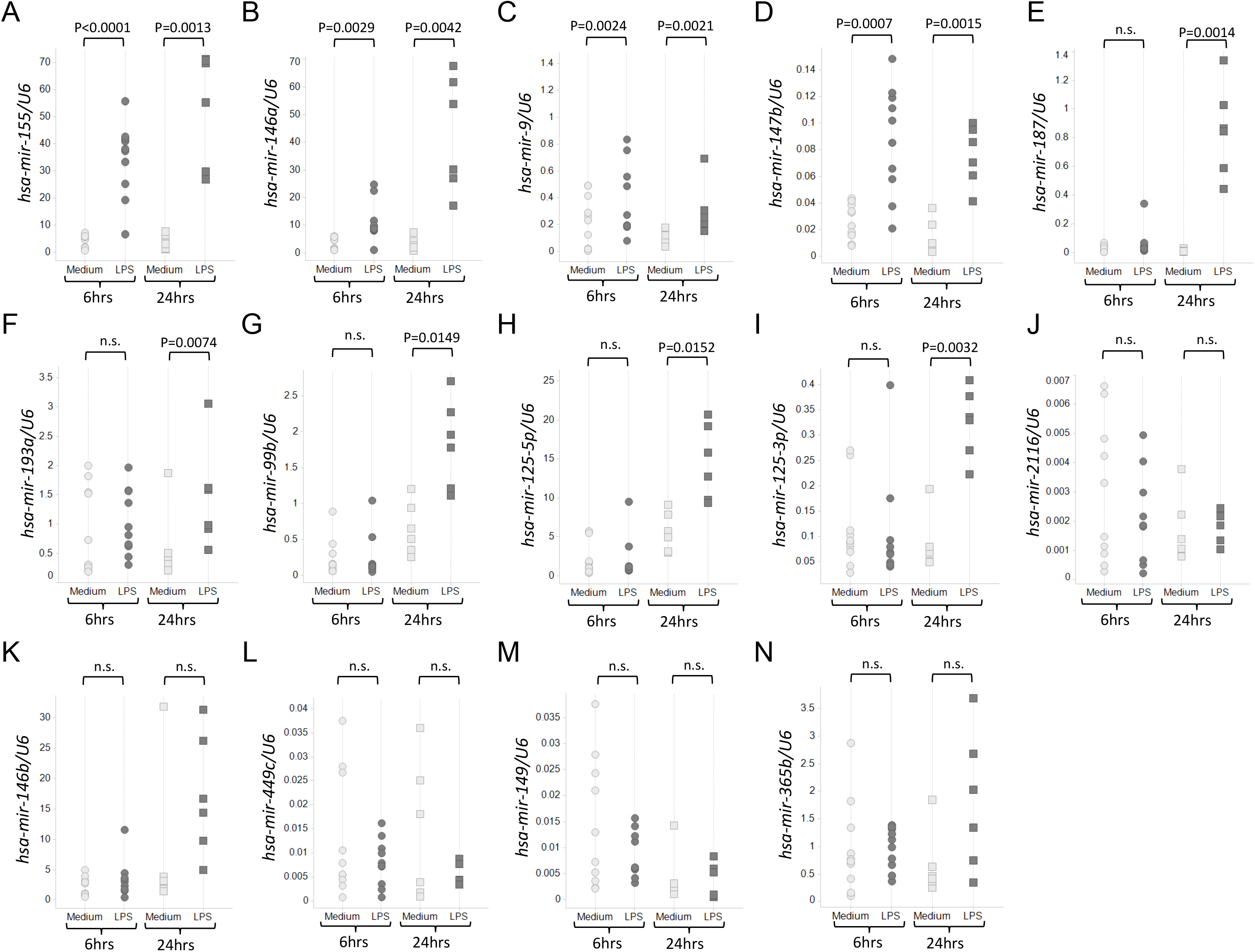
Expression validation of signature miRNAs in LPS-treated monocytes by quantitative RT-PCR. The abundance of signature miRNAs was quantified relative to U6 RNA and shown in black points for the LPS-treated monocytes or gray points for the medium-treated monocytes. Samples from nine healthy donors were tested for the 6-hour timepoint, and six healthy donors were tested for the 24-hour timepoint. Taqman-based qRT-PCR assays to detect mature miRNA expression confirmed statistically significant upregulation of *hsa-mir-155* (A), *hsa-mir-146a* (B), *hsa-mir-9* (C) and *has-mir-147b* (D) at both timepoints. qRT-PCR confirmed statistically significant upregulation of *hsa-mir-187* (E), *hsa-mir-193a* (F), *hsa-mir-99b* (G), *hsa-mir-125-5p*, (H), and hsa-mir-125-3p (I) at 24 hours. MiRNA species (J) hsa-mir-2116 (K) hsa-mir-146b, (L) *hsa-mir-449c*, (M) *hsa-mir-149*, and (N) *hsa-mir-365b* were not differentially expressed at either timepoint. P-values were determined using the two-tailed Student’s paired t-test.

To determine whether miRNAs were upregulated due to increased transcription of pri-miRNAs or due to increased pre-miRNA processing by Dicer, we tested whether we could detect upregulation of pre-cursor miRNAs in the miRNAs upregulated by LPS stimulation after 6 hours. We developed a semi-quantitative RT-PCR method to simultaneously visualize the differential expression of both mature and precursor forms of miRNA based on the differences in amplicon size. As detailed in Figure 3A, we introduced linker RNA oligos at both ends of RNA transcripts and followed this by RT-PCR with a pair of oligos located on a universal linker sequence and a miRNA-specific region. Amplified products were then resolved by gel electrophoresis, enabling separation of the smaller mature miRNA products separated from larger pre-cursor miRNA products. The assay revealed that expression of both mature and precursor forms of *hsa-mir-155*, has-*mir-146a, and has-mir-193a* were robustly induced in LPS-treated human monocytes (Fig 3B). This revealed that increased transcription of these pri-miRNAs contributed to their upregulation. LPS upregulation of the mature forms of *hsa-mir-9* and *hsa-mir-147b* could be easily detected, but we only visualized trace amounts of their precursors (Fig 3B). Such upregulation of *has-mir-9-1* precursor, which is one of pri-miRNA for production of mature *has-mir-9*, was further confirmed by using the qRT-PCR assay (Table 3). *Hsa-mir-21* expression, which were expressed abundantly in all NGS samples and served as a positive control for expressed small RNAs in human monocytes, showed even expression in mature and precursor forms regardless of LPS stimulation (Fig 3B).

**Figure 3.**
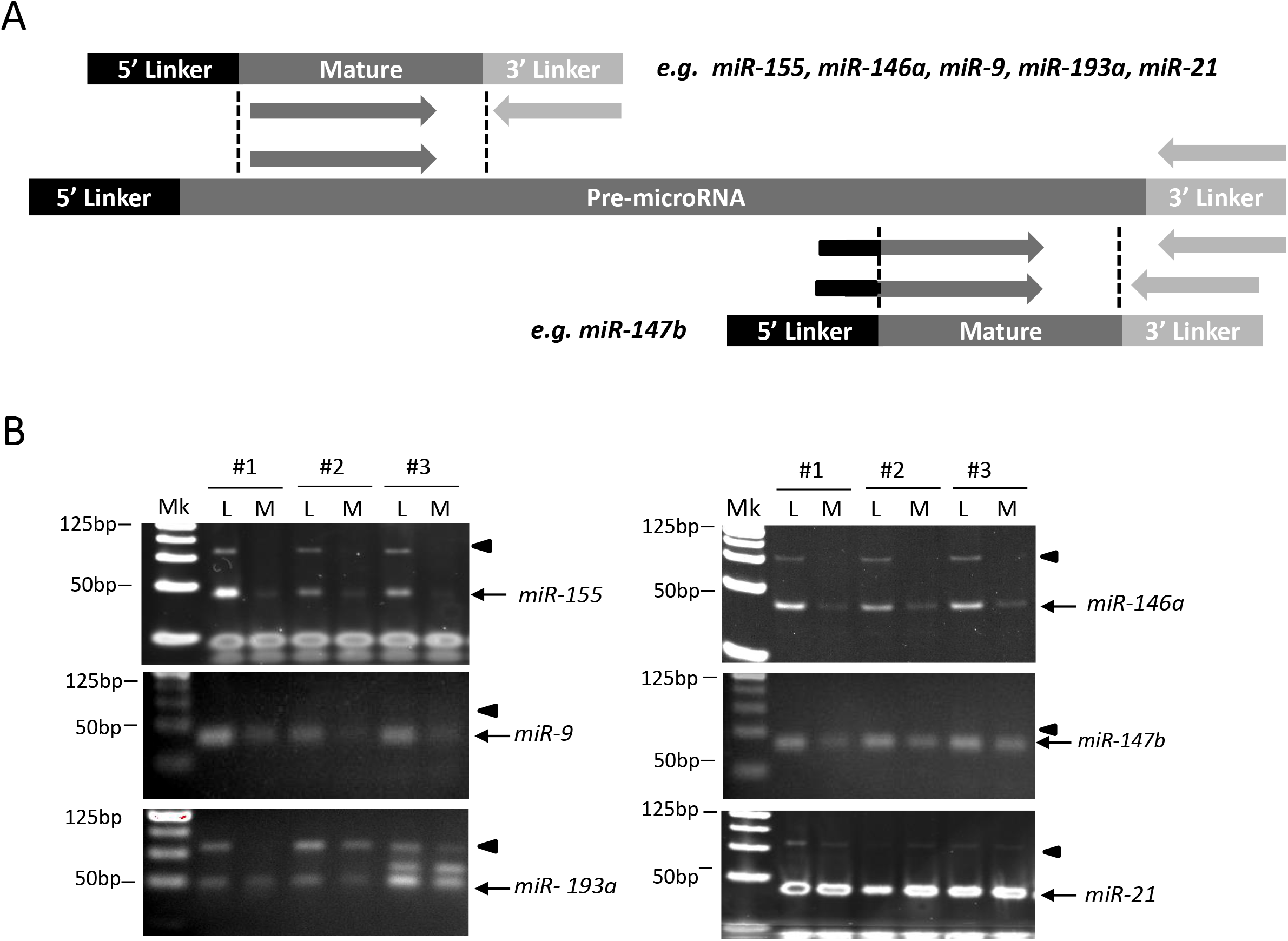
Validation of mature and precursor miRNA activation by the tagged RT-PCR assay. (A) Schematic of the tagged semi-quantitative RT-PCR strategy used for precursor and mature miRNA amplification in monocytes treated with LPS or medium alone. (B) RT-PCR analysis reveals detection of the precursor and mature forms of LPS-upregulated miRNAs *hsa-mir-155*, *hsa-mir-146a*, hsa-mir-9, hsa-mir-147b, and *hsa-mir-193a*, as well as control *hsa-mir-21* after 6 hours of culture (n=3). Black triangles denote gel band size of precursor miRNAs, while black arrows denote the mature miRNAs. Mk=marker, L=LPS-treated, M=medium-only

Overall, using a combination of small RNA-sequencing, qPCR, and RT-PCR, we identified a signature of five mature miRNAs (*hsa-mir-155, hsa-mir-146a, hsa-mir-9, hsa-mir-193a*, and *hsa-mir-147b*) whose upregulation in LPS-stimulated monocytes were validated by at least two different methods at both 6 hours and 24 hours post-stimulation (Table 3).

### Relationship between upregulated miRNAs and target mRNA expression

Having defined five signature miRNAs that were stably upregulated at 6 and 24 hours after stimulation, we next investigated how upregulation of these miRNA species affects the differential expression of target mRNA genes. We performed RNA-sequencing of 12 samples (6 LPS-treated, 6 media-treated) from 6 donors stimulated with LPS for 6 hours. We also performed RNA-sequencing of 6 samples (3 LPS-treated, 3 media-treated) from 3 donors stimulated with LPS for 24 hours. Reads were aligned to the human genome (build hg38) and DESeq2 was used to test for differential expression. In total, 734 differentially expressed genes were identified at 6 hours (462 upregulated, 272 downregulated) (Table 4) and 588 DEGs were identified at 24 hours (516 upregulated, 72 downregulated) (Table 5). 1270 unique genes were differentially expressed in total, with 52 of those genes shared at both timepoints. Principal components analysis (PCA) of all differentially expressed genes revealed that samples treated with LPS clearly clustered separately from unstimulated samples, as explained by principal component 1 (Fig 4A-B). To a lesser extent, samples treated for 6 and 24 hours also clustered separately from each other (Fig 4A-B). The differences in differentially expressed genes and global expression profiles as analysed by PCA reveal that mRNA expression in monocytes varies highly between 6 and 24 hours even though the miRNA expression profile remains relatively similar.

**Figure 4.**
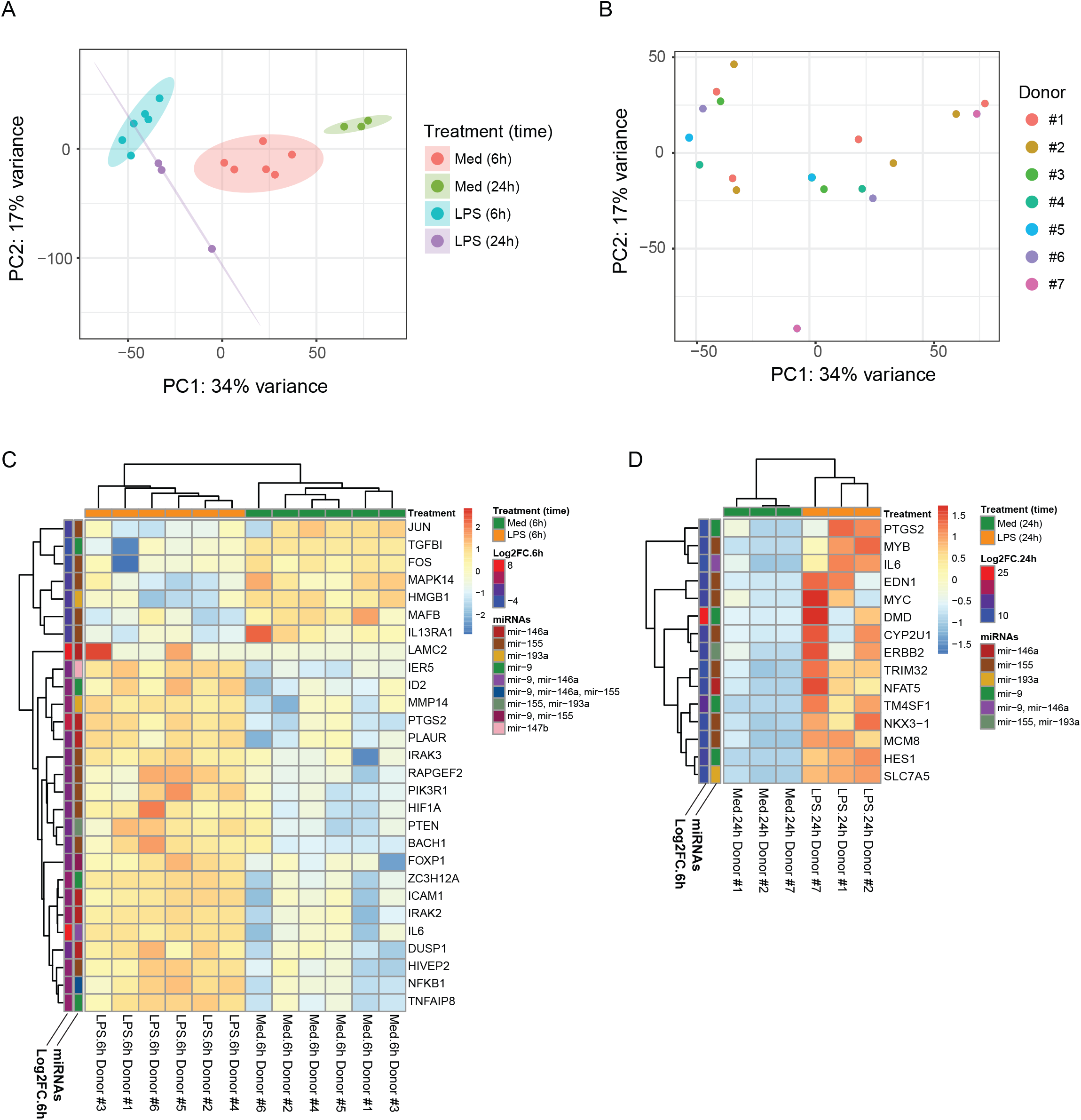
Expression analysis of previously validated miRNA targets by RNA-seq. (A and B) Principal components analysis (PCA) plot of all differentially expressed RNA-seq genes comparing LPS- and medium-treated monocytes at 6 hours and 24 hours. Samples colored by treatment and length of treatment in (A). Samples colored by sample donor in (B). (C) Z-score expression heatmap of validated miRNA targets that are also differentially expressed after 6 hours of LPS treatment. (D) Z-score expression heatmap of validated miRNA targets that are also differentially expressed after 24 hours of LPS treatment. Log2FC.6h/Log2FC.24h = log2-fold expression change of LPS-treated vs medium-only monocytes at 6 or 24 hours; miRNAs = miRNAs known to regulate given genes.

Analysis of gene ontology (GO) terms associated with the differentially expressed genes identified response to LPS, response to molecule of bacterial origin, and response to IL-1 as primarily cell functions associated with LPS stimulation at 6 hours (Supplementary Fig 5). Conversely, GO terms plasma lipoprotein particle clearance, regulation of phospholipase A2 activity, and positive regulation of myeloid cell differentiation were most significantly enriched in the genes associated with LPS downregulation at 6 hours (Supplementary Fig 5). At the 24-hour timepoint, GO terms cell migration and locomotion were associated with genes enriched by LPS stimulation, while GO terms regulation of innate immune response and humoral immune response were associated with genes downregulated by LPS stimulation (Supplementary Fig 5).

We first identified differentially expressed genes that were also functionally validated miRNA targets by miRTarBase and asked how expression of these genes varied between monocytes treated with and without LPS. At 6 hours, we identified 28 such genes (Fig 4C). Surprisingly, we found that only 7 genes (all of which are regulated by *hsa-mir-155*, except *TGFBI* and *HMGB1*, which are regulated by *hsa-mir-9* and *hsa-mir-193a*, respectively) were downregulated in LPS-stimulated monocytes, while the remaining 21 genes were upregulated in LPS-stimulated monocytes. Four genes were regulated by multiple signature miRNAs (Fig 4C). At 24 hours, we identified 15 differentially expressed genes that are also known targets for the upregulated miRNAs, of which two genes were regulated by multiple signature miRNAs (Fig 4D). All 15 genes were upregulated in LPS-stimulated monocytes, indicating a consistent pattern of gene upregulation in LPS-treated monocytes despite concurrent upregulation of miRNAs.

We also determined whether miRNA upregulation might result in lower target mRNA expression over time. Using MirTarBase, we first compiled a list of functionally validated miRNA targets genes of *hsa-mir-155, hsa-mir-146a, hsa-mir-193a, hsa-mir-9*, and *hsa-mir-147b*. Then we analysed their fold-change in expression relative to LPS at both 6 and 24 hours (Fig 5A). We identified a heterogenous pattern of target RNA expression, as some target mRNAs (e.g. *IL6*, *TNF*) had greater relative expression at 6 hours than after 24 hours, while other mRNAs (e.g. *IRAK3*, *NFKB1*) had greater relative expression after 24 hours. A slight majority of miRNA-regulated genes had greater relative expression at 24 hours relative to at 6 hours despite consistent miRNA expression at both time points (Fig 5B-F). The upregulation of these genes suggests that despite the consistency of miRNA upregulation in response to acute LPS stimulation, other factors besides miRNA likely play a major role in regulating RNA expression. The results also suggest that miRNAs in LPS-stimulated monocytes do not completely abrogate mRNA expression but may fine-tune mRNA expression to so that cellular responses can be enacted with precision.

**Figure 5.**
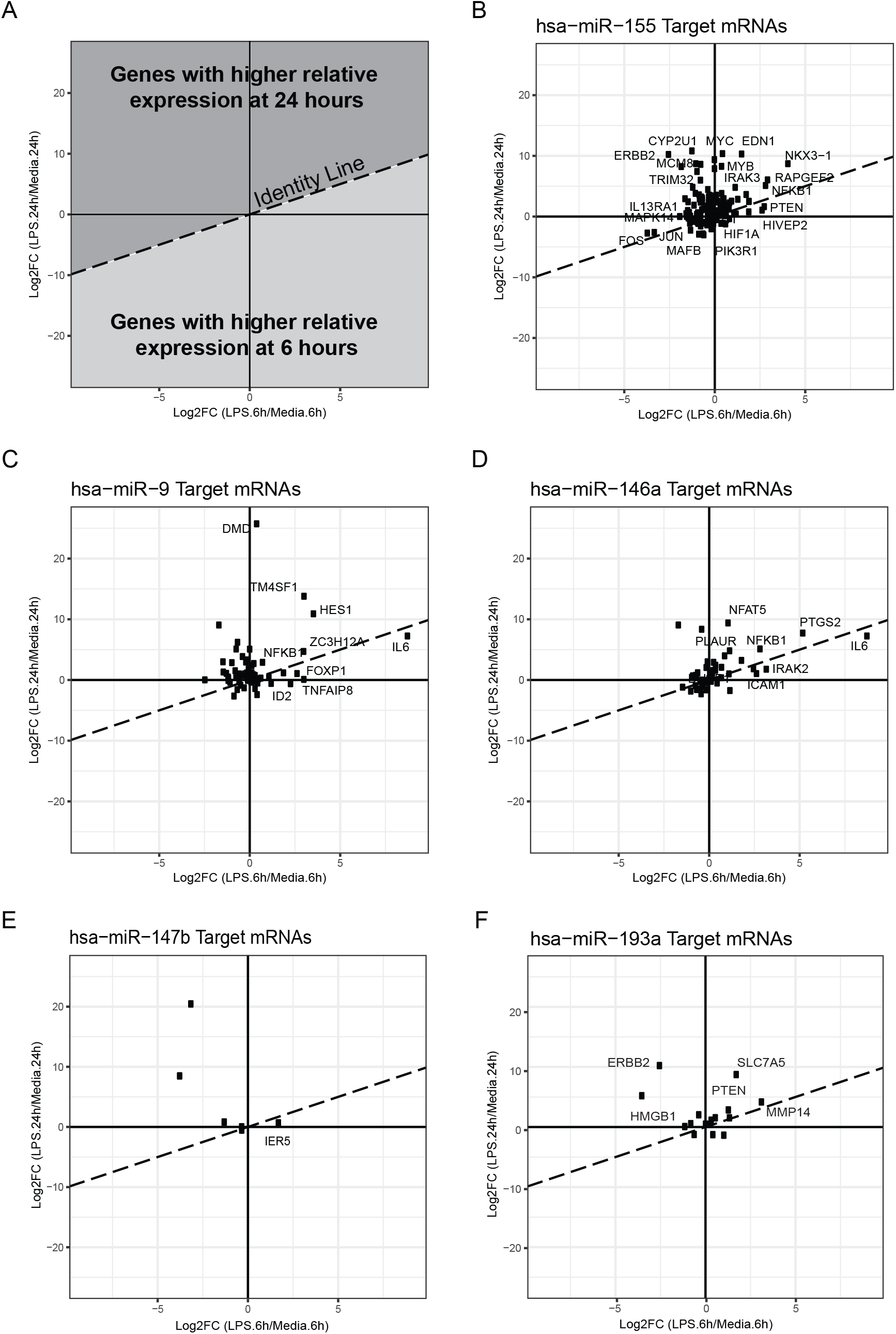
Target mRNA expression dynamics of miRNAs upregulated by LPS treatment. (A) Diagram depicting the log-fold change expression of miRNA-regulated genes after 6 hours and 24 hours of LPS stimulation. (B-E) Log-fold change RNA-seq gene expression at 6 and 24 hours of previously validated targets of miRNAs *hsa-mir-155* (B), *hsa-mir-9* (C), *hsa-mir-146a* (D), and *hsa-mir-147b* (E), and *hsa-mir-193a* (F). Black dots represent individual genes that are targets of a specific miRNA. Significantly differentially expressed genes are labeled. Dotted line represents the unity line.

### Identification of tRF RNAs derived from transfer RNA in human monocytes

In addition to the miRNAs sequences we characterized, a significant portion of the NGS reads were mapped to other small RNA species with known functions in altering cellular function. While we did not detect significant differential expression in any of the small RNA species after LPS stimulation (Supplementary Figs 6 and 7), we investigated whether any tRNA-derived fragment (tRF) RNAs were conserved in human monocytes. tRFs are derived from transfer RNAs, and several species have been cloned and identified in few organisms and cell types [16,17]. To investigate tRF RNAs in human monocytes throughout the genome, we mapped all the small RNA sequence reads systematically to the tRNA genomic database. We compared 30 bp of the flanking regions upstream or downstream to all of the 625 known or predicted tRNA gene loci that resulted from the predictions of the program tRNAscan-SE [20,21]. By screening the sequencing reads along with these tRNA regions, we generated a collection of the sequences that were 20-30 bp in size, appeared at least 500 counts per million reads and presented at least 100 times more than the 2^nd^ highest sequence in the same tRNA gene locus. This enabled us to identify stable tRF sequences that were frequently generated in monocytes after RNase Z or Dicer processing in the gene silencing complex. The enrichment threshold served to prevent detection of tRNA pieces that resulted from random degradation. Our screening strategy identified 18 tRF conserved sequences in human monocytes. These 18 tRFS could potentially be derived from a list of 50 tRNA loci that encoded 18 types of tRNA for 12 specific amino acids. Of these 18 tRFs, 7 were mapped to the 5’-end, 8 were located on the 3’-end, 1 was located in the middle, and 2 were found at the 3’ downstream of tRNA genes (Table 6). Lee *et al.* identified 17 tRFs in human prostate cancer cell lines, including 5 at the 5’-end, 8 at the 3’- end, and 4 in the downstream of the tRNA genes [24]. Three of them, *tRF-1001* (*tRF-SerTGA*), *tRF-3006* (*tRF-GlyGCC*), and *tRF-5005* (*tRF-GlyGCC*), were also found in the list of our human monocyte samples. The other 15 tRFS in our study are novel.

We performed stem loop RT-PCR analysis to detect two novel tRFs, *tRF-GlnTTG* and *tRF-TyrGTA*, and one known one*, tRF-1001*, as the control. As shown in Fig 6A, the stem loop RT-PCR assays confirmed expression of *tRF-GlnTTG*, *tRF-TyrGTA*, and *tRF-1001* in human monocytes and a higher abundance of *tRF-TyrGTA* in two LPS-treated monocyte samples than their matched controls. RT-PCR also confirmed these results for monocytes cultured for 6 and 24 hours (Fig 6B).

**Figure 6.**
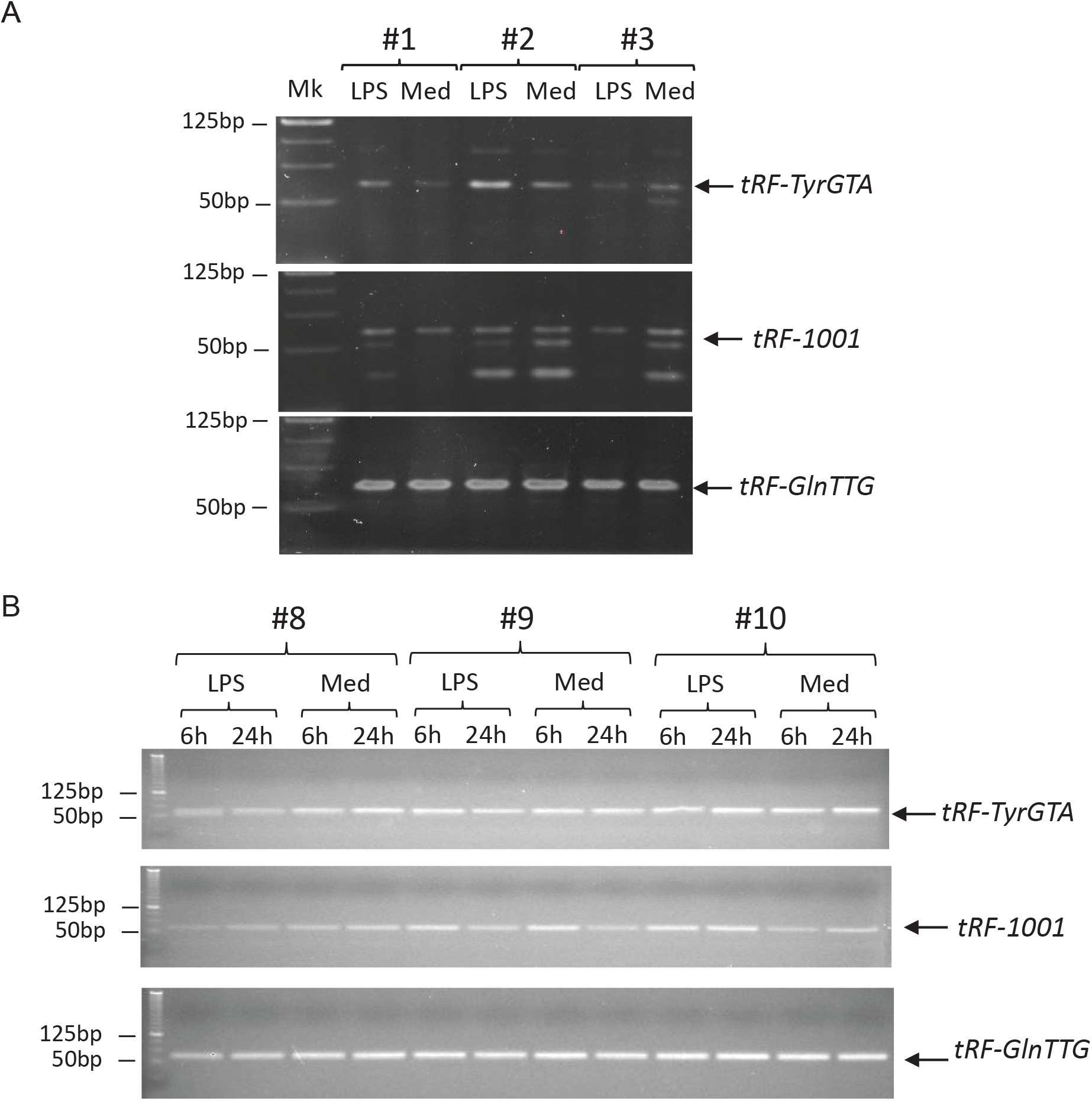
Validation of expressed tRF RNAs in human monocytes by RT-PCR. (A) The tRF products generated by the stem loop RT-PCR assays for *tRF-GlnTTG*, *tRF-TyrGTA*, and *tRF-1001* were ~57bp in size and shown as the arrow lines. (B) RT-PCR assays to validate the expression of identified tRFs at 6 and 24 hours after LPS stimulation or culture in medium only.

## DISCUSSION

In this study, we carried out a comprehensive small RNA transcriptome analysis in human monocytes from multiple donors and investigated how monocytes utilize miRNAs to regulate their response to over time during acute inflammation induced by LPS. Using NGS enabled us to globally profile all small miRNAs in an unbiased manner, allowing us to validate new miRNAs and filter out low quality sequences that are likely to be false positives. Based on the frequency of the sequencing reads mapped to human genome, we found that 72.1-95.8% of the sequencing reads were mapped to miRNAs followed by the reads mapped to rRNAs, snoRNAs, tRNAs, and hY RNAs, suggesting that miRNAs are the predominant class of small RNA species from 18-nt to 30-nt in length. The analysis also revealed disproportionate miRNA expression in monocytes, as the top hundred species of miRNAs accounted for over 90% of the miRNA population in human monocytes although over eighteen hundred miRNAs have been published in miRBase. We also identify 18 tRFs that have not been previously described in monocytes, and future studies investigating these genes will elucidate their cellular functions in monocytes. In addition to discovering miRNAs that are uniquely upregulated at 6 hours and 24 hours post-stimulation, we identified a signature of five miRNAs that were stably upregulated at both phases of inflammation. This concise list of signature miRNAs suggests that monocytes selectively activate only a few miRNA species rather than respond with a universal shift in all miRNA populations within the short-term response to LPS stimulation.

Previous investigations have used oligo- or qPCR-based arrays to identify miRNAs in myeloid cells that rapidly increase expression through activation of TLRs or TNFα (Supplementary Table 1). Most of these studies have focused on myeloid cell lines or macrophages, which differ in their gene regulation circuitry. As a result, LPS stimulation may cause slightly different miRNAs to be enriched in each cell type (Supplementary Table 1) even before accounting for differences in dosage and length of stimulation. This illustrates the importance of considering cell type, species of origin, and treatment duration when understanding miRNA expression.

Despite differences in cell type, classic miRNAs *hsa-mir-155* and *hsa-mir-146* were consistently implicated in nearly all previous investigations including our study, suggesting that these miRNAs are critical for promoting pan-myeloid function in response to LPS [9,10,25]. Moreover, since monocytes act as first responders to inflammation and can differentiate into macrophages and dendritic cells, the upregulation of both miRNAs suggests that different stages of the innate myeloid response share some similar regulatory mechanisms. *Hsa-mir-155* had been demonstrated to be involved in differentiation/proliferation of B-cells, cancers, cardiovascular disease, apoptosis, and viral infection previously [25]. Its direct targets include genes involved in transcription regulation, membrane receptors, kinases, binding proteins, and nuclear proteins [26]. *hsa-mir-155* was also shown to directly target some of the genes involved in tumor suppression and apoptosis, including *FAs-associated via death domain (FADD), Jumonji AT-rich interactive domain 2 (JARID2*), and *Src homology 2-containing inositol phosphatase-1 (SHIP1)* [26] and, also, suppressed *Caspase 1 (CASP1)* in dendritic cells [9] and *tumor protein 53-induced nuclear protein 1 (TP53INP1*) in pancreatic tumors [27]. The findings supported that *hsa-mir-155* may be implicated in anti-apoptosis or cell survival besides activation of the innate immune response*. Hsa-mir-146a* has been reported to modulate inflammatory responses in human CD14^+^CD16^−^ monocytes by inhibiting NF-kB [28].

Another study investigating *hsa-mir-146a* control of NF-kB activity also found that *hsa-mir-146a* significantly decreases IRAK1 and TRAF6 levels [29]. Both *hsa-mir-155* and *hsa-mir-146a* inhibit key signalling mediators of the TLR pathway and inflammatory response. Our results not only confirm these findings but demonstrate that they maintain consistent upregulation even 24 hours after LPS stimulation. According to the result generated by the tagged RT-PCR assays, which can detect specific miRNA sequences at all stages of cytoplasmic miRNA biogenesis, both mature and precursor *hsa-mir-155* or *hsa-mir-146* were robustly induced in LPS-treated human monocytes. This reveals that activation of these classic miRNAs occurs before the precursor miRNAs are processed, likely through transcriptional activation in the cell nucleus.

*Hsa-mir-9*, *hsa-mir-147b*, and *hsa-mir-193a* appear to be more specific to monocytes than to macrophages and other monocyte derivatives [9–11,30–36]. A previous study used Taqman Low Density Arrays to study miRNAs in human monocytes and identified *hsa-mir-9* in monocytes stimulated by LPS for up to 8 hours [11]. *Hsa-mir-9*, like *hsa-mir-146a* and *hsa-mir-155*, also directly suppresses NFKB, but uniquely interacts with IRAK2, a signalling protein that interacts with TRAF6, MYD88, and Mal to induce TLR4-mediated NFBB activation [35]. Although there is no previously published literature related to *hsa-mir-147b* in human monocytes, murine *mir-147*, a murine analogue of *hsa-mir-147b*, was reported to be induced after the stimulation of Toll-like receptors. This induction occurred though TLR4 by LPS in mouse macrophages and negatively regulated TLR-associated signalling through the suppression of IL-6 or TNFα to prevent excessive macrophage inflammatory responses [32]. Future studies will investigate whether *hsa-mir-147b* also prevents excessive monocyte-mediated inflammation and proliferation in humans. *Hsa-mir-193a*, has been reported in multiple organs to act as a tumor suppressor [36]. Although its upregulation has not been previously reported in human monocytes, its ability to bind DNA chaperone *Hmgb1*, which can bind to TLR4, suggests that *hsa-mir-193a* may help regulate excessive TLR signalling [37,38].

Bazzoni and colleagues also report that *hsa-mir-99b*, *hsa-mir-125a* and *hsa-mir-187* are induced after 8 hours of LPS treatment[11], while our study finds that these miRNA genes are upregulated after 24 hours but not after 6 hours of LPS treatment. Two possible reasons may explain why our findings differ slightly from those reported in that study. First, it is possible that these three miRNAs are induced after 8 hours but not after 6 hours of LPS stimulation. Second, RPMI1640, which was used in their study to culture cells, contains glutathione, which may exogenously prime the induction of miRNA genes in response to inflammatory stress, while X-Vivo media used in our study does not contain glutathione [39]. Regardless of the exact timepoint for activation, both studies confirm that these miRNA species become upregulated in response to LPS.

As shown in Table 7, although multiple TLR signalling pathways can induce the miRNAs we detected, the promoter regions of murine *mir-147* and human *hsa-mir-155*, *hsa-mir-146a*, and at least one of *hsa-mir-9* family members, *hsa-mir-9-1*, contained the NF-κB or AP-1 binding sites that could be responsive to LPS induction and reach maximal level within 8 hours from the activation of NF-κB or C-JUN through TLR4 signalling during pro-inflammatory immune response [9–11,31,32,40,41]. Several reports related to target identification by performing miRNA-mRNA expression profile pairing, *in silico* prediction of the target sites, and target validation *in vitro* revealed that some components or their binding partners in the MyD88-dependent proinflammatory signalling pathway appeared in the direct target list of these miRNAs. *IRAK-1* and *TRAF6* were the direct targets of both *hsa-mir-146a* [10] and *hsa-mir-147b* [32], and *NF-κB* was directly suppressed by *hsa-mir-9* [11]. *hsa-mir-155* could directly negatively modulate expression level of *IKKε*, *RIPK1*, *FOS*, *FADD*, as well as *TAB2* and *Pellino-1* that were parts of theTRAF6 complex in TLR/IL-1 signalling [9,11,30]. Although only mild suppression of these targets could be achieved by an individual miRNA, by orchestrating modulation of the key components by different miRNAs in the same cascade, we hypothesize that the subset of signature miRNAs provided monocytes an internal mechanism of actions to maintain the normal physiological function and avoid hyper-reaction of proinflammatory immune response. We propose that they do this by fine-tuning gene expression of the key components in the pathway under the impact of LPS stimulation, while exposing bacterial antigens to macrophages or other antigen presenting cells (Fig 7). However, this hypothesis does not rule out the other possible functions of these miRNAs in monocytes since dozens of the targets, validated or predicted, were still found outside of TLR/IL-1 or TNF? pathways [25,30,42].

**Figure 7.**
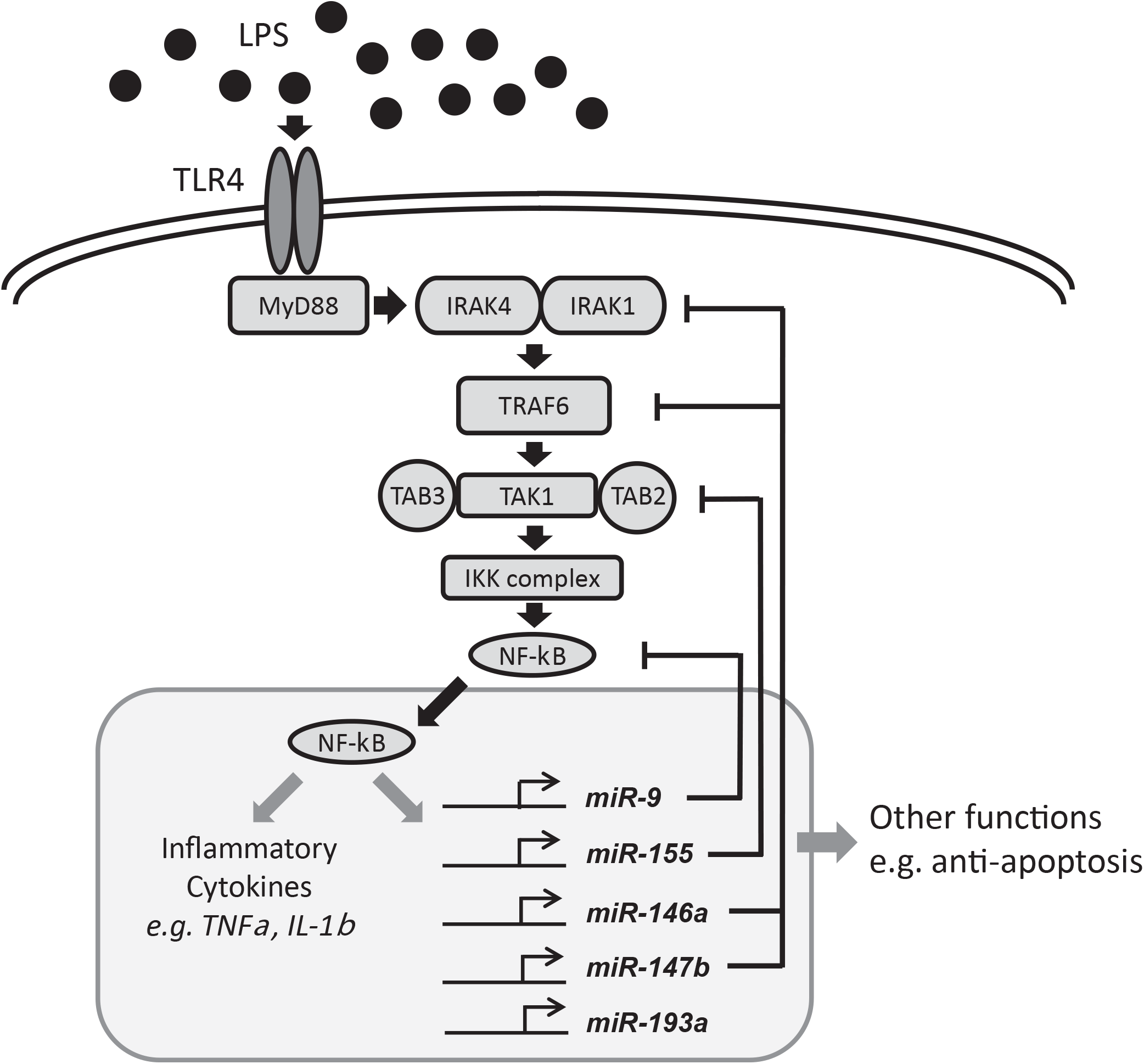
The negative feedback of signature miRNAs in the MyD88-dependent proinflammatory signalling in monocytes. Upon activation by LPS, TLR4 dimer binds to an adaptor protein, e.g. MyD88, via the cytoplasmic domain. MyD88 in turn recruits IRAK1 and IRAK4 and, subsequently, IRAK4 activates IRAK1 by phosphorylation. After phosphorylation, both IRAK1 and IRAK4 dissociate from the MyD88-TLR4 complex and associate temporarily with TRAF6 leading to its ubiquitination. Following ubiquitination, TRAF6 forms a complex with TAB2/TAB3/TAK1 to induce TAK1 activation. TAK1 then associates to the IKK complex leading to IκB phosphorylation and, subsequently, nuclear localization of NF-κB. Activation of NF-κB triggers the production of inflammatory cytokines such as TNF-α, IL-1 and IL-12, and, also, up-regulation of signature miRNAs, including *hsa-mir-155*, *hsa-mir-146a*, *hsa-mir-147b*, and *hsa-mir-9*. In the negative feedback loop, *IRAK-1* and *TRAF6* are direct targets for both *hsa-mir-146a* and *hsa-mir-147b. NF-kB* is directly suppressed by *hsa-mir-9*, and *hsa-mir-155* could directly negatively modulate the expression level of *IKKe* and *TAB2* and *Pellino-1* which are part of the TRAF6 complex in TLR/IL-1 signalling. TNF expression is suppressed by *hsa-mir-187*.

We also investigated the impact of upregulated signature miRNAs on target mRNA expression. Although miRNAs post-transcriptionally inhibit target mRNAs and can mediate mRNA degradation in minutes [43], we find that the overall abundance of target mRNAs depends on the specific gene. For example, while many genes regulated by upregulated miRNAs showed a decrease in overall expression from 6 hours to 24 hours, over 50% of target genes had higher relative abundance after 24 hours relative to 6 hours. This demonstrates the complex relationship between miRNA and mRNA, as the rate of new mRNA transcription and miRNA-mediated decay can oscillate over time. Since numerous other mechanisms, such as transcription factor recruitment, transcription initiation, and P-body activity, all affect mRNA levels, our results suggest that miRNAs play a role in fine-tuning mRNA responses rather than drastically influencing absolute expression levels. Such activities enable cell responses to be rapidly and precisely tuned to exogenous and internal cellular stimuli.

At the end of the study, we identified and reported a subset of small RNAs, tRFs, that were frequently mapped to the distinct portions of several specific tRNAs. Under our screening criteria, 18 tRF conserved sequences were identified according to the public tRNA database. Three of our 18 tRFs were published as *tRF-1001* (*tRF-SerTGA*), *tRF-3006* (*tRF-GlyGCC*), and *tRF-5005* (*tRF-GlyGCC*) by *Lee et al.* in human prostate cancer cells, LNCap, and PC-3 [24], and also detected in HEK293s, Hela cells, or a nasopharyngeal carcinoma cell line, 5-8F. In addition, a 5’ tRF from *tRNA-LeuCAG* and the 3’ trailer tRF from *tRNA-ThrAGT* in our list were identified in HEK293 cells and in 5-8F cells, respectively [44]. However, the physiological functions of tRF small RNAs had not been well defined, and only limited information related to tRFs was available in the published literature. Of the known tRF small RNAs, it has been reported that tRF-1001 was highly expressed in several carcinoma cell lines [24]. According to recent studies, mainly done in HCT-116 cells, actions related to gene silencing by some of tRF small RNAs have been investigated and detected. Haussecker *et al*. showed that, with involvement of Argonaut proteins (AGO), the tRNA-derived small RNAs, e.g. *cand14* and *cand45* (a.k.a. *tRF-1001*), were able to suppress the activity of luciferase protein expressed by the report construct containing a complementary tRF target sequence at the 3’-downstream of the luciferase ORF cDNA in dual luciferase assays [45]. By introducing the siRNA oligos against *tRF-1001* into HCT-116 cells, Lee *et al.* identified an inhibitory effect of tRF-1001 siRNA oligos on cell proliferation that could be rescued by the addition of wild-type *tRF-1001* oligos [24]. Along with identification of novel small RNA sequences (Data not shown), our study was the first systematic study to investigate and detect tRF RNA species in a subset of human immune cells by NGS and, then, verified by stem loop RT-PCR analysis, potentially leading to an interesting field in investigating tRF functions in the immune cells.

## CONCLUSION

By deep sequencing, we have highlighted a subset of signature miRNAs that are up-regulated and proposed to be implicated in the negative-feedback control of proinflammatory response in human monocytes after 6 hours and 24 hours of LPS stimulation. We also found that while these miRNAs are upregulated during LPS treatment, the overall expression level of these target genes in RNAs is still higher in LPS-treated monocytes, suggesting that miRNA inhibition may subtly modulate expression levels rather than completely halt expression. We also provided the first evidence of novel tRF RNA expression in human monocytes. In addition, our study provides a global view of small RNA transcriptome in human monocytes that agrees with and supports previous findings in individual miRNAs or tRF RNAs. Meanwhile, our study leads us in new directions and suggests further experiments that will help us to understand how these small RNAs regulate and modulate gene expression and cellular function in the innate immune system.

## COMPETING INTERESTS

The authors have read the journal’s policy and have the following conflicts: Chi-Ming Li, Daniel Lu, Tracy Yamawaki, Hong Zhou, Edwin Lamas, and Songli Wang are employees at Amgen Inc. Wen-Yu Chou is a contract worker for Amgen Research. Sabine S. Escobar, Heather A. Arnett, Huanying Ge, and Mark Chhoa were employees of Amgen while working on the study. All of the authors owned Amgen shares when the experiments were carried out. However, these do not alter the authors’ adherence to all the journal policies on sharing data and material.

## AUTHOR CONTRIBUTIONS

CML initiated, designed, and carried out the experiments and authored the manuscripts. DL, TY, and HG performed data analysis and coauthored the manuscript. EL and WC carried out the NGS run and coauthored the section of methods for the manuscript. HZ and MC carried out the experiments for expression validation. SSE and HAA prepared the monocyte samples for the experiment and critically evaluated the manuscript. DL, SW, and CML made contributions to experiment design and advised critically on data analysis, validation, and manuscript preparation. All authors read and approved the final draft of the manuscript.

## ACKNOWLEDGEMENTS

We would like to thank Chris Lindvay and Amgen NGS wet-lab team for the technical support and Robert Sandrock and Todd Juan for helpful suggestions in experiment design. This study was funded and supported by Amgen Inc., which played a role in study design, data collection and analysis, decision to publish, and preparation of the manuscript.

## ETHICS STATEMENT

The samples were collected anonymously in an internal blood donor program that was monitored by the Western IRB board in Amgen Washington. The data in this manuscript were analyzed anonymously and not linked to any identifying personal information. Thus, the study was considered exempt from IRB approval.

---

**Supplementary Figure 1.**
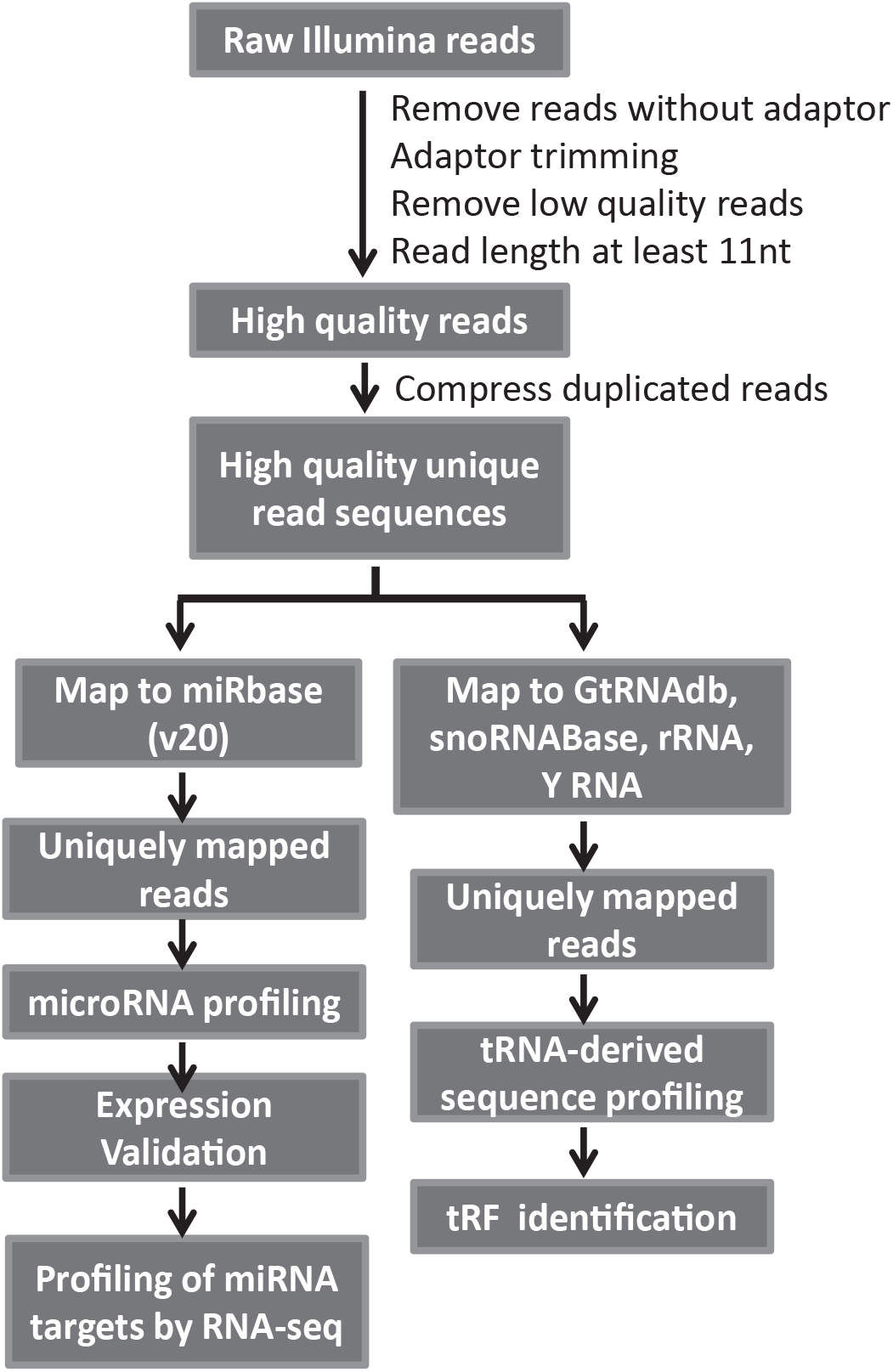
Analysis pipeline for the identification of small RNAs in expressed human monocytes stimulated by LPS. Low quality sequencing reads were first filtered by removing low quality reads (Phred score<30 or no sequencing adaptor), removing read fragments less than 12 nucleotides in length, and trimming adaptor sequences flanking the reads. Unique sequences were then aligned either miRbase (v20) and other small RNA reference databases that included human tRNA sequences from GtRNAdb, snoRNA/scaRNA sequences from snoRNABase, as well as ribosomal RNAs and hY RNAs 1/3/4/5 from NCBI, using OmicSoft Aligner (V5.1). Multi-mapped reads were discarded while uniquely mapped reads were retained for downstream small RNA identification, profiling, and validation.

**Supplementary Figure 2.**
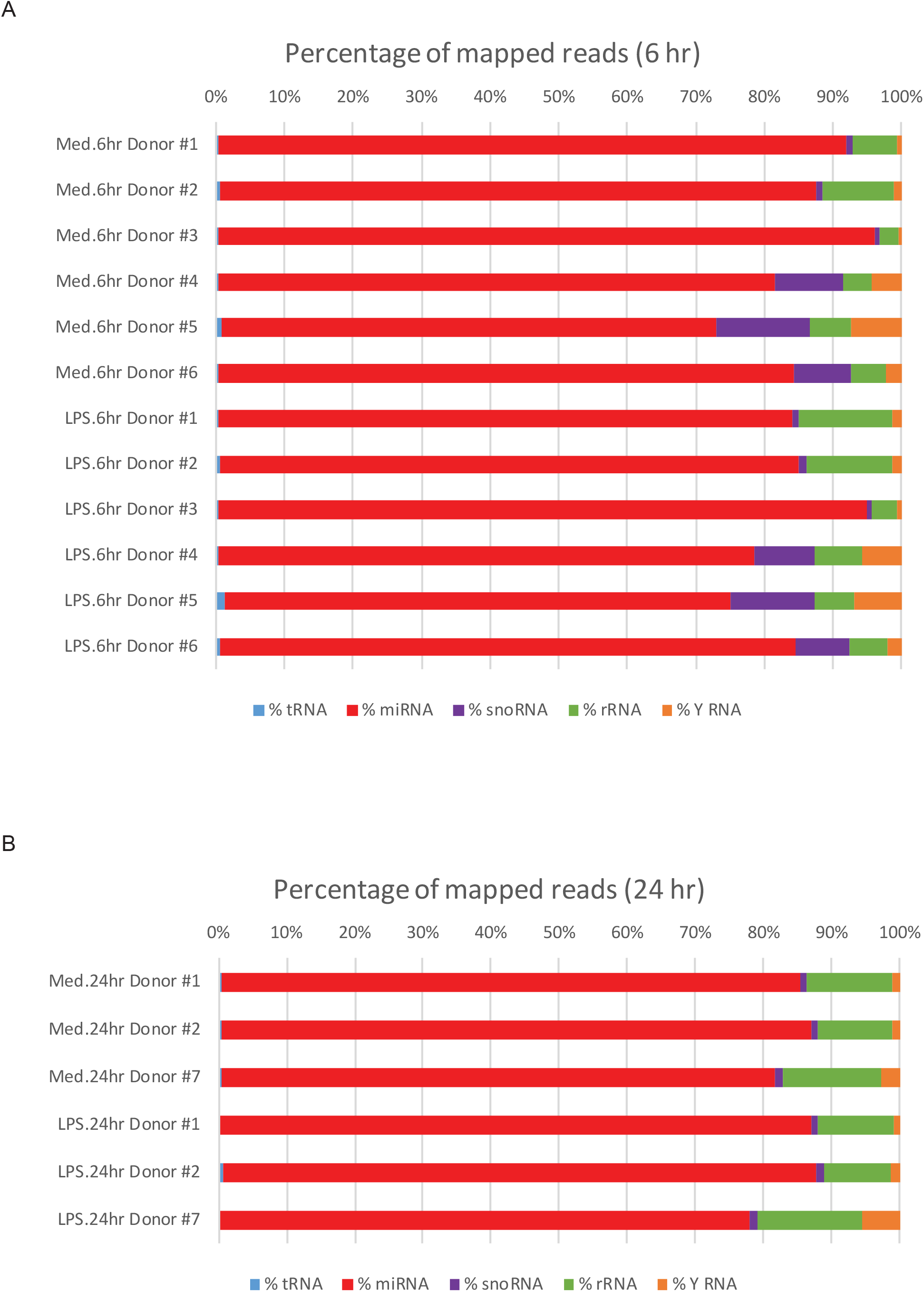
Mapping summary of small RNA transcriptome analysis in human monocytes. The percentage of reads aligned to the various RNA species are indicated in the bar plots. Blue = tRNA, red = miRNA, purple = snoRNA, green = rRNA, orange = Y-RNA. (A) shows samples sequenced after 6 hours of treatment, while (B) shows samples sequenced after 24 hours of treatment. For all samples, 72.1-95.7% of reads aligned to miRNAs.

**Supplementary Figure 3.**
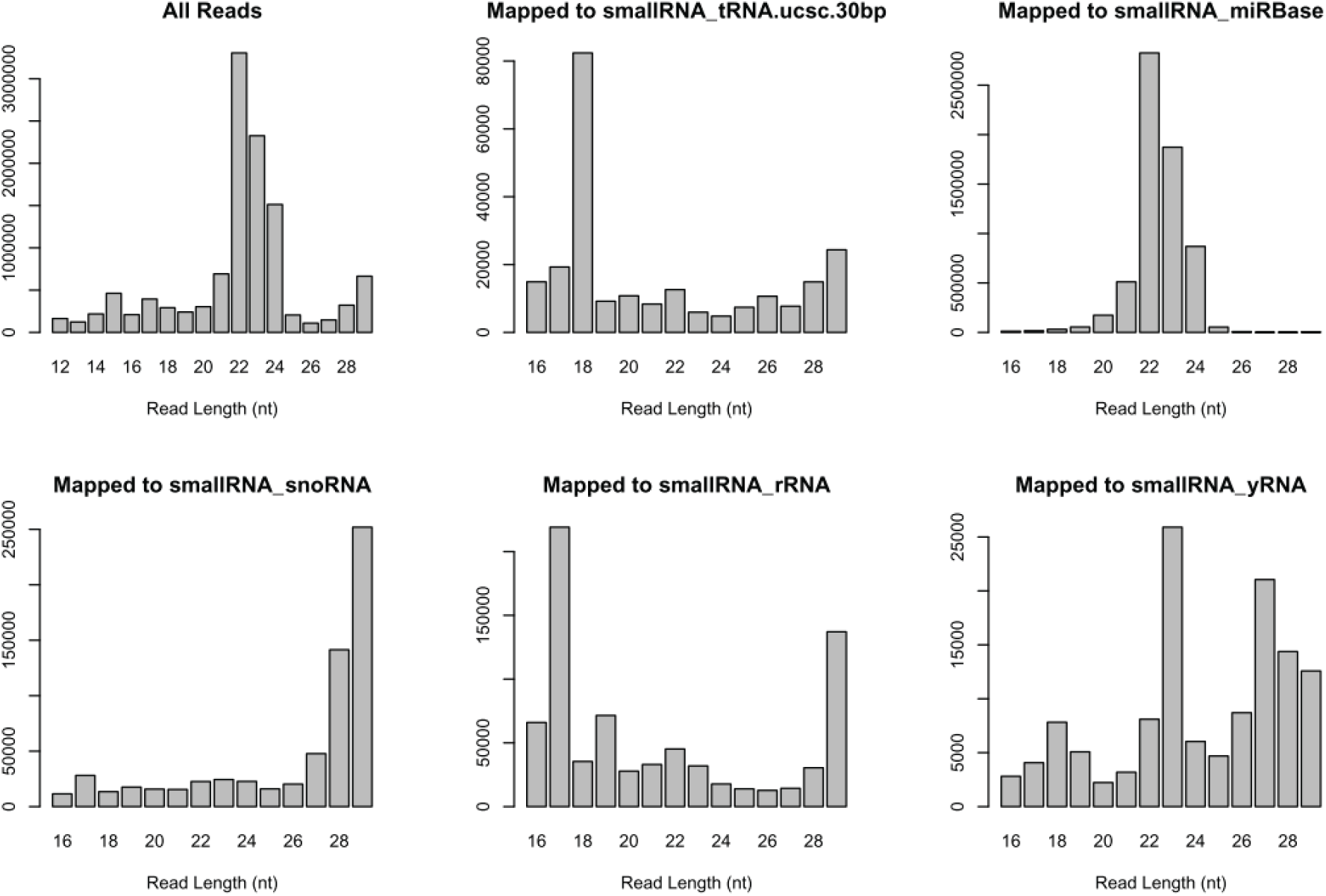
Read length distribution of mapped small RNAs. The distribution of the insert sizes of the small RNA libraries show that the insert sizes of the reads were predominantly 19bp to 24bp in length, whereas the length of the inserts for miRNA ranged from 20 to 24bp and that of tRNA-derived small RNAs was concentrated at 18bp.

**Supplementary Figure 4.**
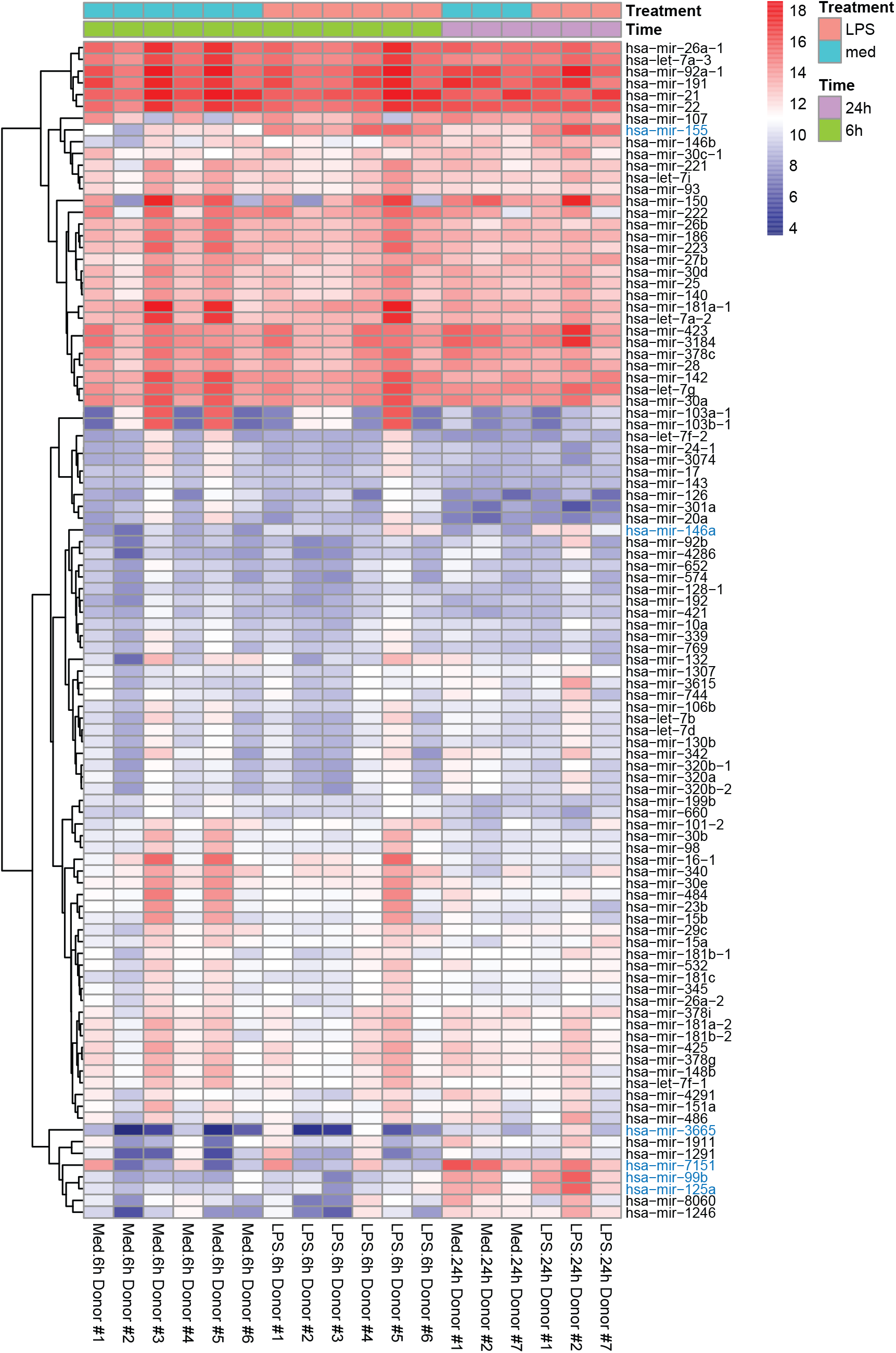
Top 100 highly expressed miRNAs in monocytes stimulated with LPS for 6 or 24 hours. Normalized expression of miRNAs is shown in log2(counts per million + 1) for each sample. Blue miRNAs denote significantly miRNAs at either 6 hours or 24 hours post-stimulation with LPS.

**Supplementary Figure 5.**
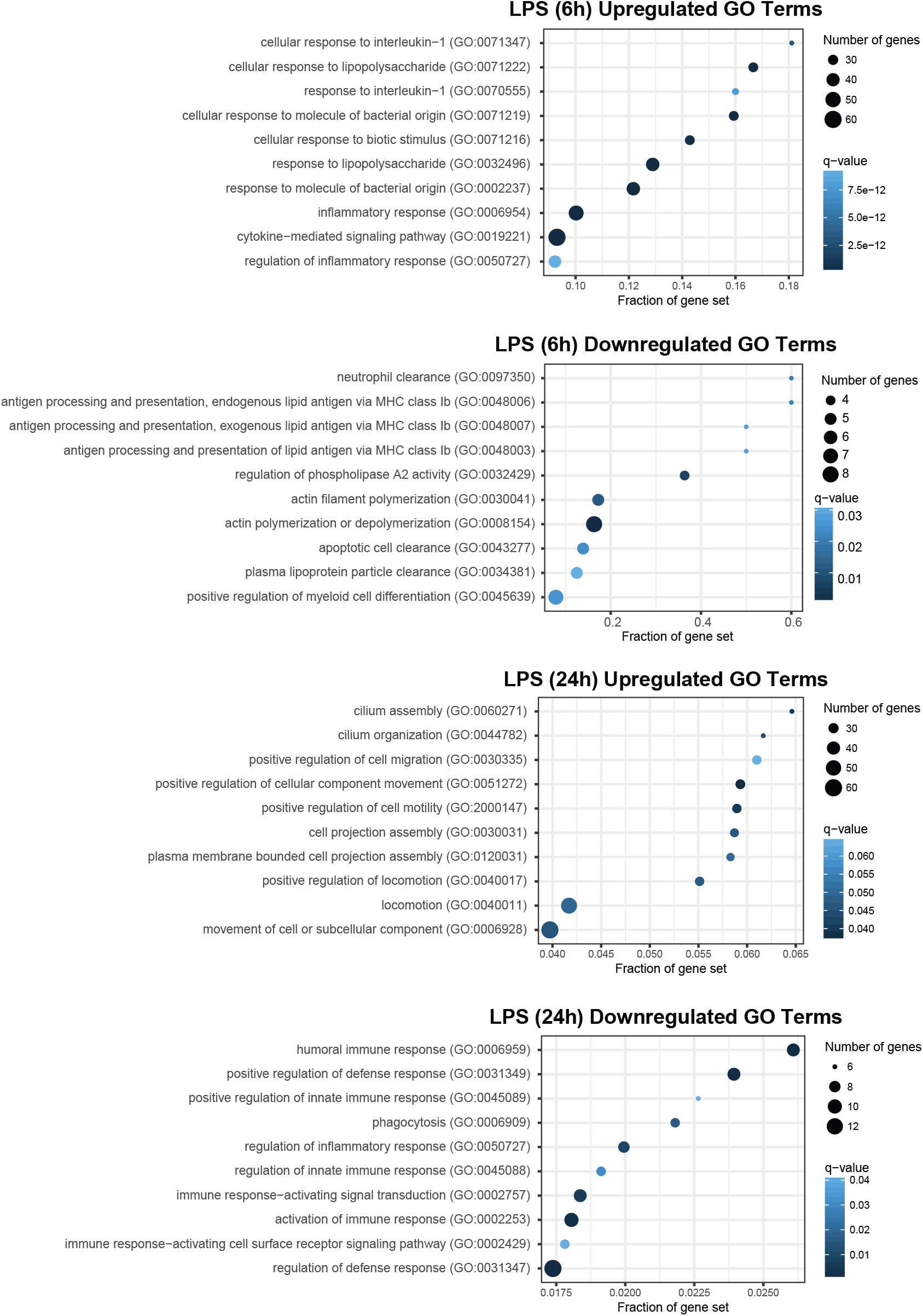
Gene Ontology (GO) terms identified by PantherDB using genes that are differentially expressed in LPS-stimulated human primary monocytes at 6 hours and 24 hours. The top ten upregulated/downregulated GO terms with the lowest q-values are shown. Dot size represents the number of differentially expressed genes present for each GO term. Dot color represents the FDR-adjusted q-value (FDR=0.01) for each GO term.

**Supplementary Figure 6.**
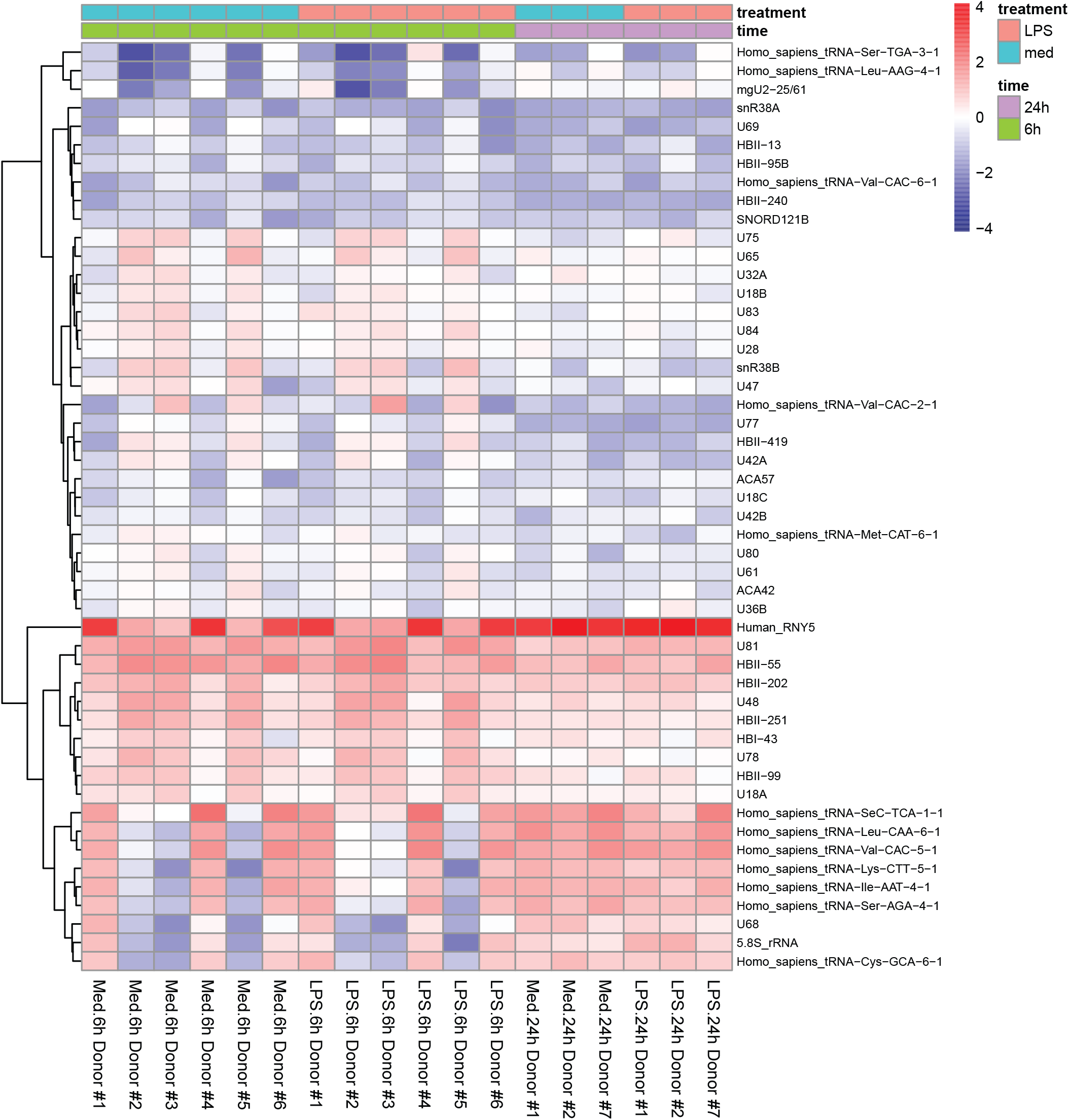
Heatmap of most variant non-miRNA small RNAs in LPS-stimulated human primary monocytes. Z-score expression heatmap of the top 50 most variant non-miRNA small RNAs in LPS-stimulated monocytes after 6 hours and 24 hours of treatment. No small RNAs had significant differential expression.

**Supplementary Figure 7.**
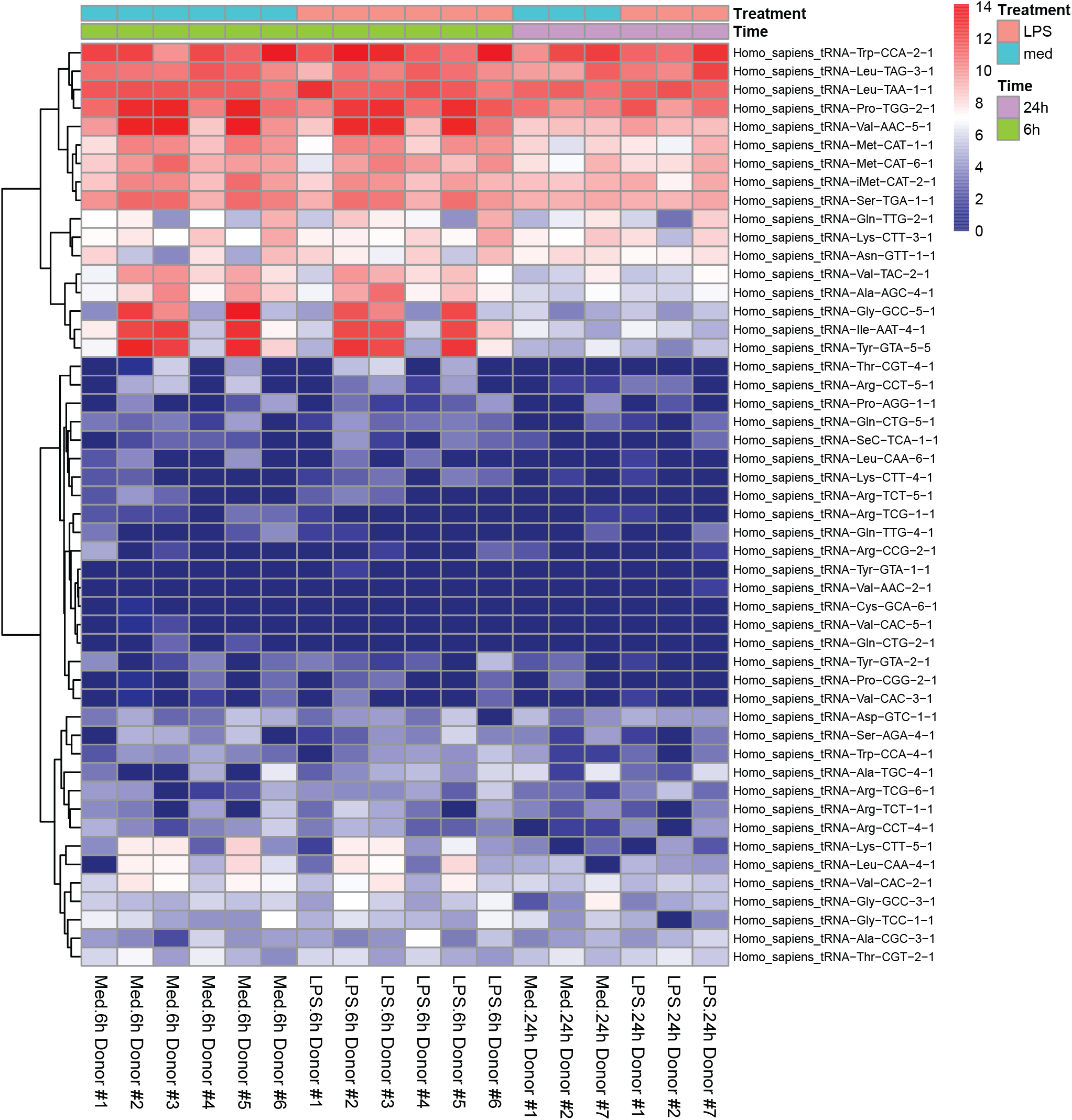
Heatmap of the most highly expressed tRNAs in LPS-stimulated human primary monocytes. Log2-fold expression heatmap of the top 50 tRNAs with the highest average small RNA-seq expression across all samples.

